# Hepatic urea cycle suppression drives brown fat thermogenesis to counter obesity and liver disease

**DOI:** 10.64898/2026.06.02.729618

**Authors:** Laura C. Kim, Nicholas P. Lesner, Xuanyan Cai, Xu Han, Jimmy P. Xu, Andy Zhao, Nathan J. Coffey, Denise Zheng, Brian Keith, Clementina Mesaros, Zoltan Arany, M. Celeste Simon

## Abstract

The hepatic urea cycle is consistently suppressed in liver disease and hepatocellular carcinoma (HCC), but whether loss of individual enzymes contributes to disease progression remains unknown. We investigated the role of urea cycle enzyme argininosuccinate synthase 1 (ASS1) in diet- and carcinogen-induced liver disease using hepatocyte-specific deletion of *Ass1* in mice. Complete loss of hepatic *Ass1* is lethal, but a high-fat diet extends lifespan. Unexpectedly, animals with approximately 85% loss of hepatic *Ass1* are markedly protected from diet-induced obesity, liver steatosis, fibrosis, and HCC. Hyperammonemia induced by hepatic *Ass1* loss activates brown adipose tissue fatty acid oxidation, driving thermogenesis and increasing energy expenditure. Importantly, targeting *Ass1* after obesity onset promotes weight loss and reverses liver steatosis. These findings implicate hepatic ASS1 as a critical regulator of systemic lipid metabolism with therapeutic potential against obesity, liver disease, and HCC.

## Main Text

The hepatic urea cycle converts toxic ammonia generated from protein catabolism into urea for excretion (**Fig. 1A**)(*1*), yet is universally suppressed in multiple cancer types including hepatocellular carcinoma (HCC) and clear cell renal carcinoma(*2–4*). In established cancer cell lines, suppression of Argininosuccinate Synthase 1 (ASS1), a key urea cycle enzyme that condenses citrulline and aspartate to form argininosuccinate, promotes tumor cell proliferation by preserving aspartate pools required for nucleotide synthesis(*5*). Ureagenesis is also impaired in patients with metabolic dysfunction-associated steatotic liver disease (MASLD) and steatohepatitis (MASH)(*6–8*). Whether suppression of individual urea cycle enzymes contributes to liver disease progression or HCC initiation remains unknown and is a question of particular importance as an increasing proportion of HCC cases are associated with MASLD and MASH(*9*).

**Figure 1.**
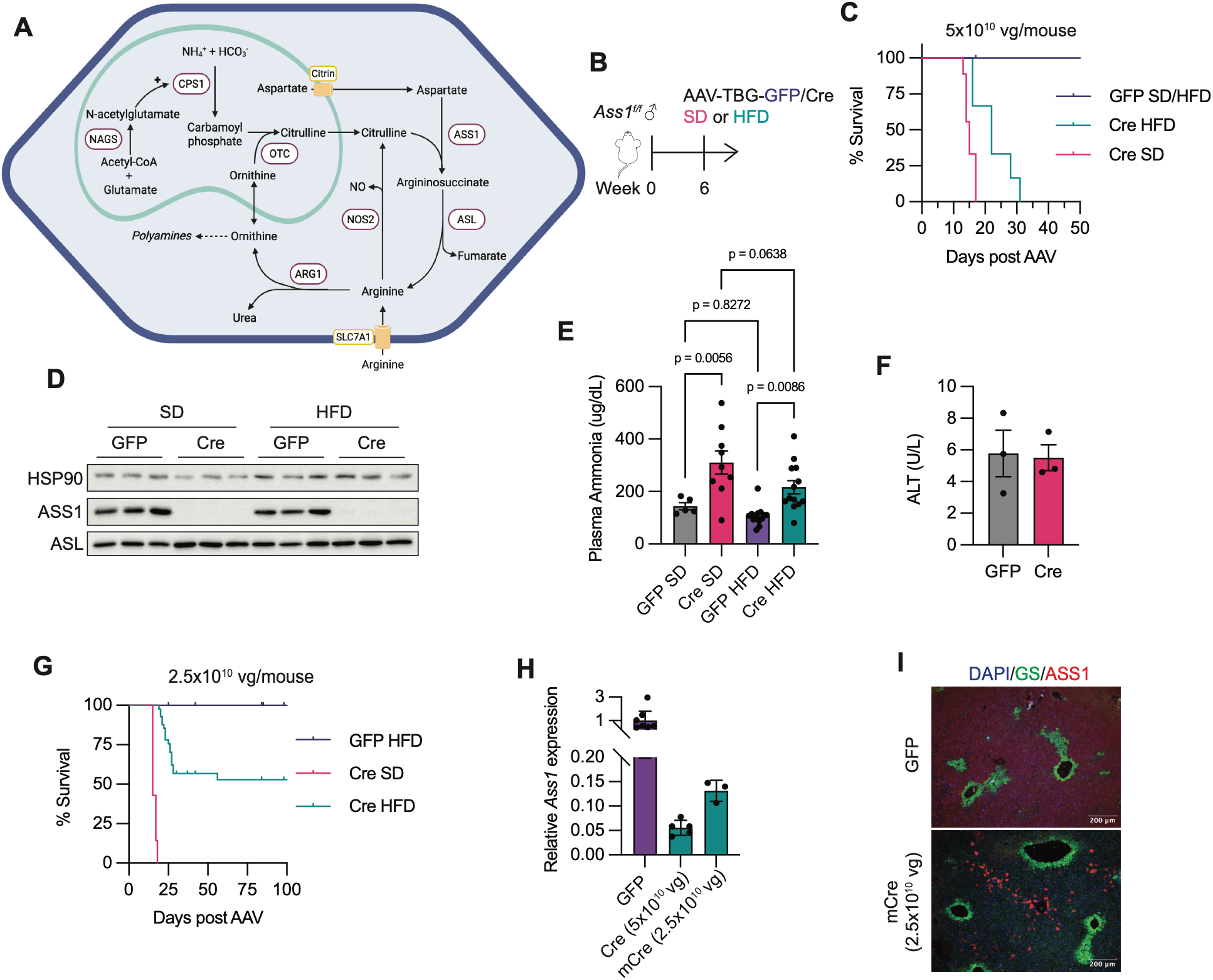
High-fat diet rescues lethality induced by loss of hepatic *Ass1*. (**A**) Schematic depicting the hepatic urea cycle. (**B)** Experimental design to test control (GFP) and hepatocyte-specific ASS1 knockout (Cre) mice on standard (SD) or high-fat diet (HFD). Argininosuccinate Synthase 1, ASS1. (**C**) Survival curve of GFP and Cre mice on SD or HFD. GFP, n=13; Cre SD, n=9; Cre HFD, n=6. (**D**) ASS1 and ASL protein expression in GFP and Cre livers on SD or HFD 13 days after AAV delivery (n=3 individual mice per group). ASL, argininosuccinate lyase; HSP90, heat shock protein 90. (**E**) Plasma ammonia levels of GFP and Cre mice on SD or HFD 12-13 days after AAV delivery. GFP SD, n=5; Cre SD, n=9; GFP HFD, n=12; Cre HFD, n=13. (**F**) Plasma ALT levels of GFP (n=3) and Cre (n=3) mice on SD 13 days after AAV delivery. (**G**) Survival analysis of GFP (n=24) and mosaic hepatocyte specific ASS1 knockout mice (mCre) on SD (n=7) or HFD (n=41). (**H**) *Ass1* mRNA expression in livers of GFP (n=7), Cre (n=5), and mCre (n=3) animals on HFD. (**I**) Representative immunofluorescence staining of ASS1, GS, and DAPI in livers of GFP and mCre mice. Glutamine Synthetase, GS.

Hypomorphic mutations in human urea cycle genes result in urea cycle disorders (UCDs). ASS1 deficiency causes Citrullinemia Type 1 (CTLN1), characterized by neonatal hyperammonemia and hypercitrullinemia that can be fatal if left untreated. Management of CTLN1 symptoms includes low-protein diet, ammonia scavengers, and arginine supplementation; however, liver transplantation remains the only curative treatment option(*10*). Guidelines for patients with UCDs include lifelong monitoring for liver diseases, such as liver failure, steatosis, fibrosis, and HCC(*11*). Whereas links between liver disease and other UCDs, particularly argininosuccinate lyase (ASL)(*12*) and citrin deficiency(*13*, *14*), are well-documented, scant evidence supports CTLN1-dependent predisposition to liver disorders and HCC(*15*, *16*). These data suggest that individual urea cycle enzymes play distinct roles in liver disease and HCC development.

Here, we show that whereas complete loss of murine hepatic *Ass1* is lethal, mosaic deletion protects mice maintained on a high-fat diet (HFD) from obesity, steatosis, fibrosis, and HCC through an unexpected systemic mechanism linking the liver urea cycle to peripheral energy expenditure. Targeting hepatic *Ass1* also reverses established obesity and liver steatosis, implicating ASS1 as a potential therapeutic target.

### High fat diet rescues lethality caused by hepatic *Ass1* loss

To explore the role of hepatic ASS1, we employed animals expressing a conditional *Ass1*^fl/fl^ allele and delivered AAV8-TBG-Cre (5×10^10^ viral genomes [vg]/mouse, “Cre”) by retro-orbital injection to specifically delete *Ass1* in hepatocytes (**Fig. 1B**). Animals injected with AAV8-TBG-GFP (GFP) served as controls. As with untreated CTLN1 infants, liver-specific *Ass1* deletion caused rapid death of animals fed a standard chow diet (SD) within approximately 14 days (**Fig. 1C**). Surprisingly, HFD (60% kcal from fat) extended lifespan despite only a 2.5% reduction in dietary protein content (**Fig. 1C, Table S1**). HFD did not alter hepatic ASS1 protein levels (**Fig. 1D**) or normalize plasma ammonia levels (**Fig. 1E**). Moreover, lethality of hepatic *Ass1* loss is unrelated to liver failure, as plasma ALT was unchanged in Cre animals (**Fig. 1F**).

Given the rapid lethality caused by complete loss of *Ass1* in the liver, we developed a mosaic model of hepatic *Ass1* loss, in which 50% of the original dose (2.5×10^10^ vg, “mCre”) of virus was administered to animals (**Fig. 1G**). mCre animals survived longer than Cre animals on SD but still died by ∼21 days, whereas more than half survived beyond 20 weeks on HFD (**Fig. 1G**). mRNA levels of hepatic *Ass1* were reduced by nearly 85% in mCre livers compared to GFP controls but were nevertheless ∼2.5-fold higher than those in Cre mice receiving the original AAV dose of 5×10^10^ vg per mouse (**Fig. 1H**). Immunofluorescent analysis of ASS1 protein levels in these livers indicates that even with reduced AAV8-TBG-Cre doses, a vast majority of hepatocytes lose ASS1 expression (**Fig. 1I**). Heterozygous deletion of *Ass1* in hepatocytes (*Ass1*^flox/+^) is not lethal and animals were phenotypically unremarkable compared to GFP controls (**Fig. S1A-C**). Together, these data suggest that ∼15% residual hepatic ASS1 expression, combined with HFD, is sufficient to sustain life.

### Mosaic deletion of hepatic *Ass1* protects against diet-induced obesity and liver steatosis

UCDs are associated with development of liver steatosis and related diseases(*11*). Strikingly, mCre animals maintained on HFD were completely protected from weight gain over the course of 20 weeks, in contrast to GFP controls (**Fig. 2A, B**). Brown adipose tissue (BAT) and white adipose tissue depots were reduced in mCre mice, whereas quadriceps muscle mass was unchanged (**Fig. 2C**). Liver weights were normal (**Fig. 2D**); however, mild liver steatosis was evident in GFP control animals but not in mCre livers (**Fig. 2E, F**). mCre livers exhibited reduced neutral lipid accumulation (**Fig. 2G**) and triglyceride levels (**Fig. 2H**), confirming protection from hepatosteatosis. Plasma triglycerides trended higher, consistent with breakdown of adipose stores (**Fig. 2I**). At 20 weeks, neither GFP nor mCre animals exhibited markers of liver damage (**Fig. 2J, K**) despite chronic HFD feeding; however, mCre animals had elevated plasma ammonia (**Fig. 2L**) and citrulline (**Fig. 2M, S2A**) levels. Metabolomic analysis of livers revealed persistent increases in urea cycle intermediates in mCre animals as expected (**Fig. S2B, C**).

**Figure 2.**
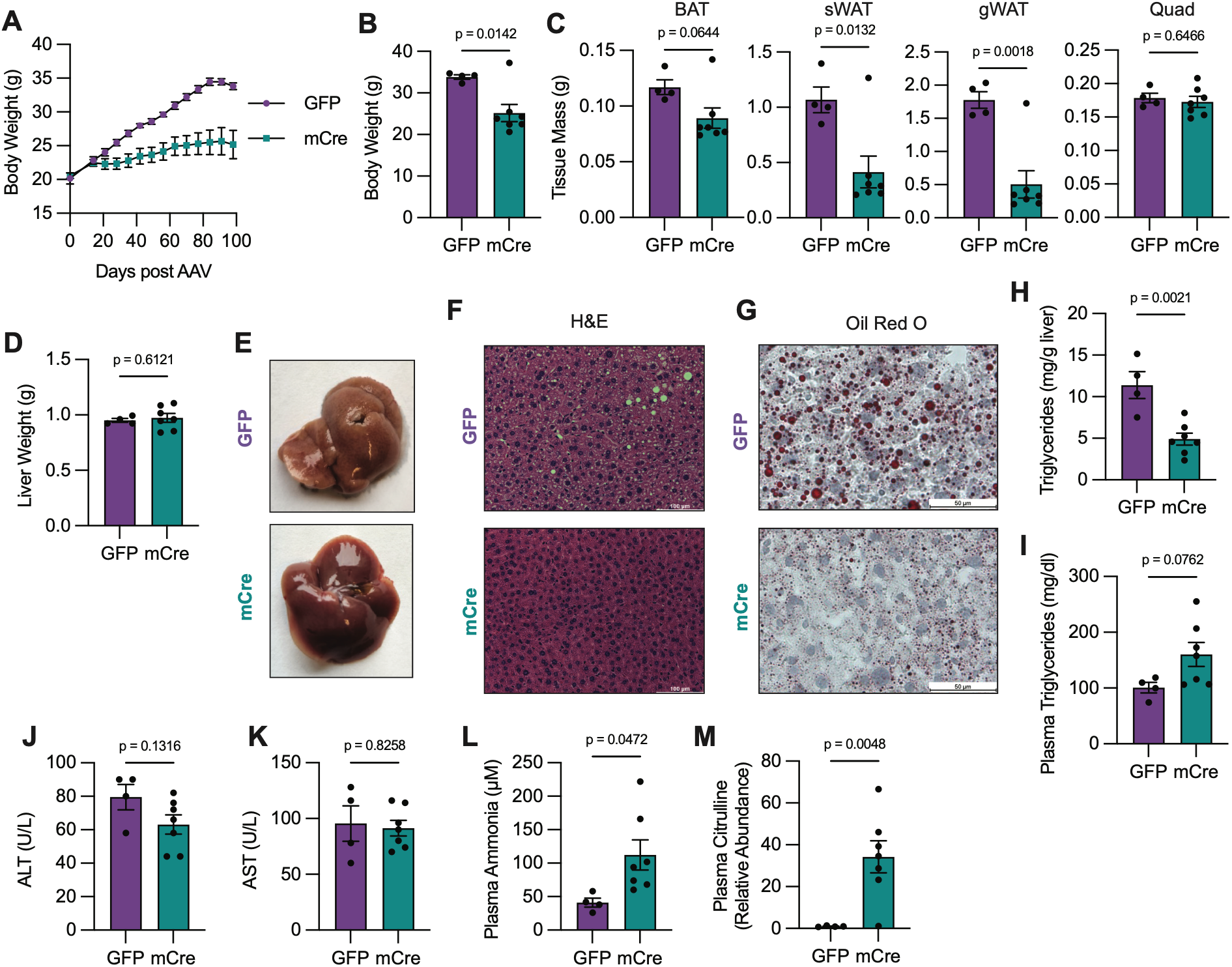
Mosaic deletion of hepatic *Ass1* protects from HFD-induced obesity and liver steatosis. (**A-L**) GFP (n=4) and mCre (n=7) animals were fed HFD for 20 weeks. (**A**) Body weights. (**B**) Body weight at sacrifice. (**C**) Tissue weights. BAT, brown adipose tissue; sWAT, subcutaneous white adipose tissue; gWAT, gonadal white adipose tissue; Quad, quadriceps. (**D**) Liver weight. (**E**) Representative images of gross livers. (**F**) Representative images of liver H&E staining. (**G**) Representative images of liver Oil Red O staining. (**H**) Liver triglycerides. (**I**) Plasma triglycerides. (**J**) Plasma ALT. (**K**) Plasma AST. (**L**) Plasma Ammonia. (**M**) Plasma Citrulline.

At 36 weeks on HFD, GFP animals exhibited exacerbated obesity-related phenotypes, while mCre animals remained protected from weight gain, adiposity, and steatosis (**Fig. S3A, B**). mCre animals were additionally protected from hepatomegaly and elevated fasting blood glucose levels associated with prolonged HFD feeding (**Fig. S3C-E**). A carbohydrate-free ketogenic diet (80% kcal from fat; **Table S1**) similarly extended mCre survival and reduced adiposity (**Fig. S4A-D**), demonstrating that dietary carbohydrates have no apparent effect on mCre phenotypes. Together, these data challenge the widely held assumption that *Ass1* loss increases susceptibility to hepatosteatosis, while also suggesting that a chronic HFD could benefit patients with hypomorphic *ASS1* alleles.

### Mosaic deletion of hepatic *Ass1* protects against MASH-HCC

To investigate whether *Ass1* loss also protected against other liver disease phenotypes, we fed GFP control and mCre animals the Gubra-Amylin NASH (GAN) diet for 25 weeks (**Fig. 3A**). The GAN diet (high fat, fructose, and cholesterol; **Table S1**) recapitulates human MASLD and MASH phenotypes including obesity, metabolic syndrome, liver steatosis, and fibrosis. As with HFD, mCre animals fed a GAN diet had reduced body weight (**Fig. 3B, C**), liver weight (**Fig. 3D**), and adipose depots (**Fig. 3E**). GFP animals fed GAN chow had more pronounced liver steatosis than that observed with HFD chow alone. In contrast, mCre animals were spared from developing liver steatosis (**Fig. 3F, G**) and exhibited reduced neutral lipid droplet accumulation (**Fig. 3H**). mCre animals also exhibited reduced liver fibrosis, as indicated by reduced collagen deposition (**Fig. 3I**) and fibrotic gene expression (**Fig. 3J**).

**Figure 3.**
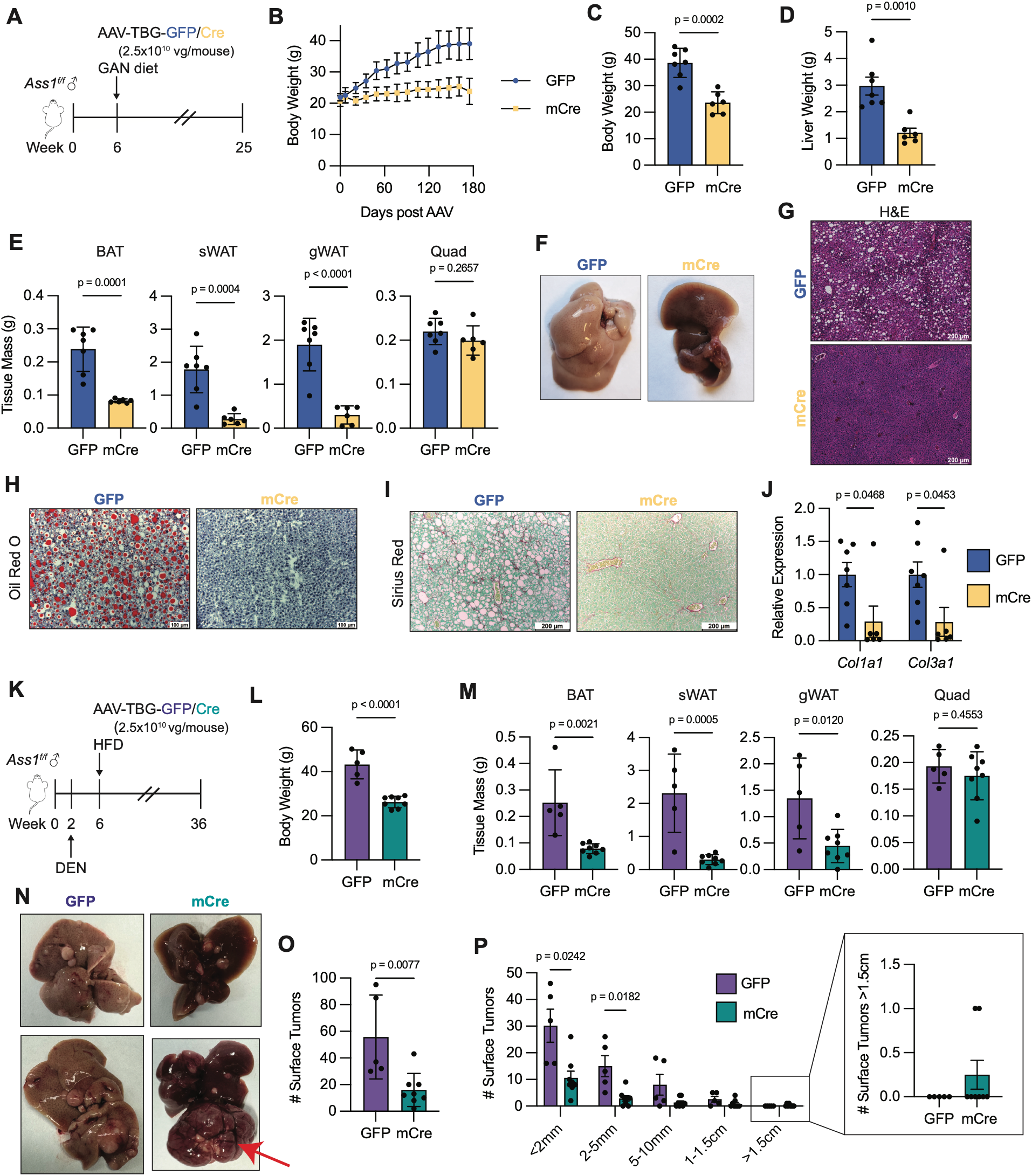
Mosaic deletion of hepatic *Ass1* protects against MASH and HCC. (**A-J**) GFP (n=7) and mCre (n=6) animals were fed the Gubra-Amylin NASH (GAN) diet for 25 weeks beginning at time of AAV delivery. (**A**) Schematic model of experimental design. (**B**) Body weight over time. (**C**) Body weight at sacrifice. (**D**) Liver weight. (**E**) Tissue weights. (**F**) Representative images of gross livers. (**G**) Representative images of liver H&E staining. (**H**) Representative images of liver Oil Red O staining. (**I**) Representative images of liver Sirius Red staining. (**J**) qRT-PCR analysis of liver fibrosis markers *Col1a1* and *Col3a1*. (**K-P)** Diethylnitrosamine (DEN, 25mg/kg) was administered i.p. to *Ass1*^fl/fl^ mice at 2 weeks of age. At 6 weeks, AAV was delivered to generate GFP (n=5) or mCre (n=8) animals and HFD feeding was started. Animals were monitored for 36 weeks, then sacrificed for analysis. (**K**) Schematic model of experimental design. (**L**) Body weight at sacrifice. (**M**) Tissue weights. (**N**) Representative images of gross livers. Arrow indicates tumor larger than 1.5cm. (**O**) Total surface tumor counts per liver. (**P**) Surface tumor size distribution. Inset enlarges surface tumor counts larger than 1.5cm.

We then investigated whether hepatic *Ass1* loss could affect spontaneous, autochthonous MASLD-HCC development using the chemical carcinogen diethyl-nitrosamine (DEN) in combination with a HFD (**Fig. 3K**). At 36 weeks, mCre animals were again protected from obesity (**Fig. 3L**) and had reduced adipose stores (**Fig. 3M**). Importantly, mCre animals exhibited reduced liver adiposity and tumor incidence (**Fig. 3N-P**). mCre animals exhibited fewer tumors across nearly all size categories, though a small number of extremely large (>1.5cm) tumors were noted (**Fig. 3N, P**).

### Mosaic deletion of hepatic *Ass1* in diet-induced obesity model restores animals to normal body weight

Considering the protective effects of *Ass1* suppression, we investigated the therapeutic potential of targeting hepatic ASS1 after disease onset. *Ass1*^fl/fl^ animals fed HFD from 6 weeks of age were randomized after 12 weeks to receive AAV8-TBG-GFP or -Cre for mosaic *Ass1* deletion, as previously described. Animals were maintained on HFD for 10 additional weeks (**Fig. 4A**). At randomization, animals averaged over 35g (**Fig. 4B**). mCre animals began losing weight approximately 3 weeks after AAV delivery and plateaued at ∼27g by week 7, whereas GFP animals continued to gain weight (**Fig. 4B, C**). Animals appeared healthy throughout the duration of the experiment despite high plasma ammonia levels induced by loss of liver *Ass1* (**Fig. 4D**). Moreover, targeting *Ass1* after obesity onset led to reductions in adipose tissue stores and prevented hepatomegaly associated with long-term HFD feeding (**Fig. 4E, F**). Importantly, liver steatosis was also reduced (**Fig. 4G, H**). Together, these data indicate that appropriately titrated hepatic *Ass1* inhibition can reverse obesity and liver steatosis, even in the context of a HFD.

**Figure 4.**
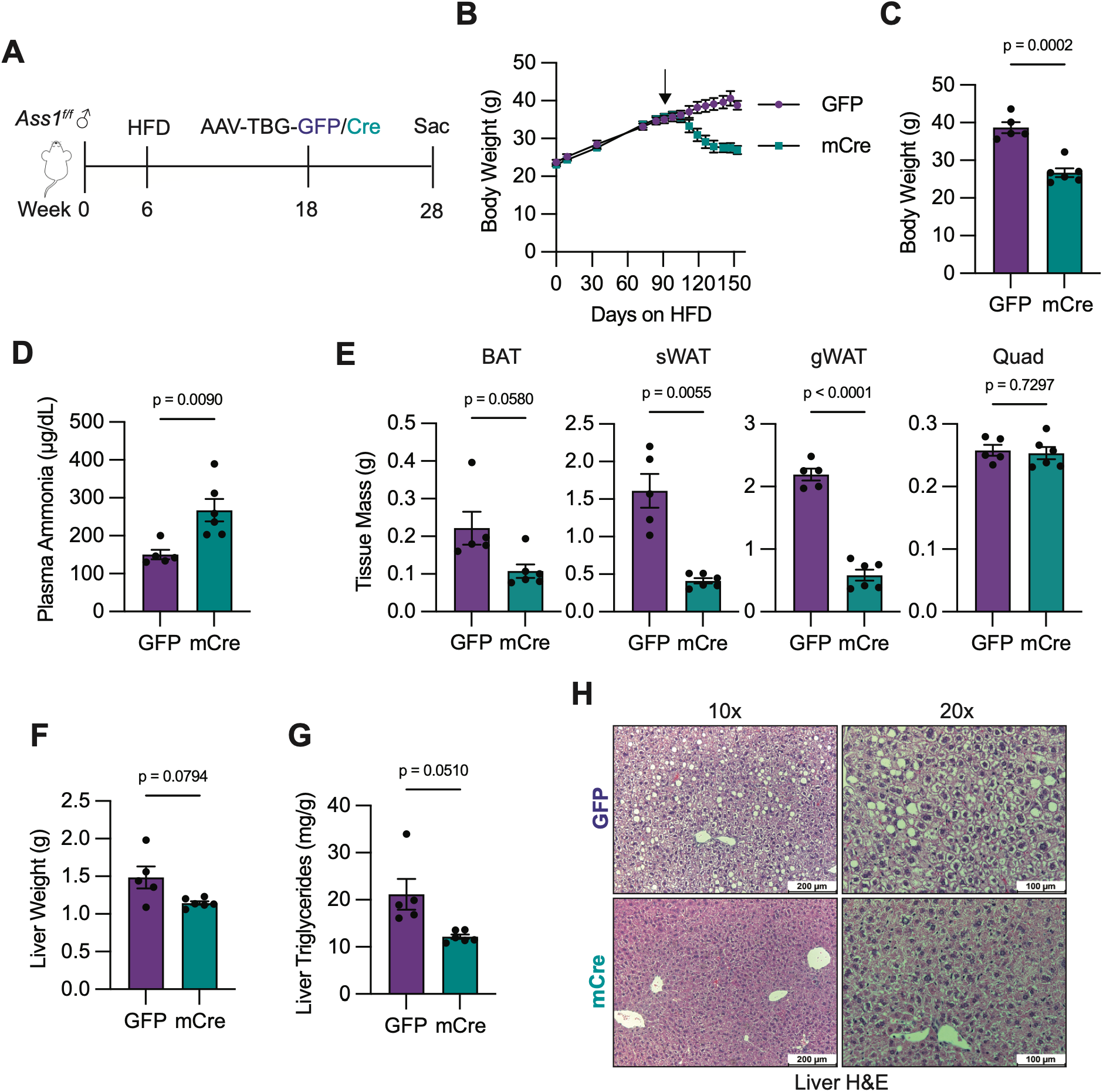
Mosaic deletion of hepatic *Ass1* promotes weight loss in obese mice. (**A-H**) *Ass1^fl/fl^* mice were fed HFD beginning at 6 weeks of age. After 12 weeks, AAV8-TBG-GFP control or -Cre (2.5×10^10^ vg/mouse) was administered and animals were monitored for 10 additional weeks until sacrifice. (**A**) Schematic model of experimental design. (**B**) Body weight. Arrow indicates time of AAV delivery. GFP n=5; Cre n=6. (**C**) Body weight at sacrifice. (**D**) Plasma ammonia at sacrifice. (**E**) Tissue weights. (**F**) Liver weight. (**G**) Liver Triglycerides. (**H**) Representative H&E images of liver tissue.

### ASS1 supports hepatic triglyceride stores

We then investigated which lipid species were specifically altered by liver *Ass1* loss by performing lipidomics on liver tissue from GFP and mCre animals 3 weeks post AAV delivery (**Fig. S5A**). Importantly, body weights and adiposity have not yet diverged at this time point (**Fig. 2A**). Overall, differences in lipids were modest, although reduced phospholipid species were noted (**Fig. S5B, C**). Quantification of individual lipid classes revealed a striking reduction of total triglyceride abundance across nearly every detectable species (**Fig. S5D, E**), indicating overall suppression of triglyceride accumulation in response to *Ass1* deletion. There were no apparent changes in the total abundance of diglycerides, lysophospholipids, or phospholipids (**Fig. S5F-H**).

To investigate mechanisms of triglyceride depletion (**Fig. S6A**), we performed bulk RNA-sequencing on livers from GFP and Cre animals maintained on a SD for 13 days or HFD for 3 weeks (**Fig. 1B**). In both cohorts, the fatty acyl-CoA biosynthesis gene set was significantly enriched in GFP samples (**Fig. S6B-E**). The most downregulated genes were related to very long chain fatty acid (VLCFA) elongation, long chain acyl-CoA synthesis, and lipid desaturation (**Fig. S6C, E**). These gene expression changes were consistent across long-term HFD (**Fig. S6F**) or GAN diets (**Fig. S6G**). Despite reproducible gene expression changes, lipidomic analysis did not reveal reductions in sphingomyelins, which contain VLCFAs as critical components (**Fig. S6H**), nor were there alterations in the lipid saturation ratio of 18:1 to 18:0 chains present in triglycerides (**Fig. S6I**). Collectively, these data suggest that hepatic *Ass1* loss disrupts liver triglyceride accumulation, although not through suppression of lipid biosynthesis.

### Liver lipid metabolism does not mediate protection from steatosis induced by loss of *Ass1*

We next tested lipid uptake, lipolysis, VLDL secretion, and fatty acid oxidation (FAO) as alternative mechanisms of liver triglyceride regulation (**Fig. S6A**)(*17*). To investigate lipid uptake, we examined mRNA expression levels of liver fatty acid transporters *Slc27a2*, *Slc27a4*, *Slc27a5*, and *Cd36*, and observed no differences between GFP and Cre livers in SD-fed animals (**Fig. S7A**). Despite the reduction in steatosis, *Slc27a5* and *CD36* were slightly upregulated in the livers of HFD fed Cre animals (**Fig. S7B**). We then measured liver free fatty acid (FFA) levels and found that livers from SD fed Cre animals had reduced FFAs, whereas HFD chow rescued FFAs at the same time point (**Fig. S7C**). Glycerol levels were reduced in Cre livers fed either SD or HFD (**Fig. S7D**), suggesting that FFA levels were maintained through uptake and not lipolysis, which would naturally generate glycerol. Thus, fatty acid transport is unimpaired and does not mediate protection from liver steatosis in mCre animals.

The liver packages excess triglycerides into very-low-density lipoprotein (VLDL) particles for secretion and delivery to adipose and skeletal muscle tissues for storage or energy(*18*). High rates of VLDL secretion can therefore deplete the liver of its triglyceride stores and prevent steatosis (**Fig. S6A**). However, VLDL secretion rates were unchanged in Cre and mCre animals compared to GFP controls (**Fig. S7E, F**).

Finally, we compared hepatic FAO rates in isolated hepatocytes from HFD-fed GFP or Cre animals. FAO rates (**Fig. S7G, H**), plasma ketone levels (**Fig. S7I**), and liver acyl-carnitine levels (**Fig. S7J**) were comparable between GFP and Cre animals, consistent with unchanged rates of hepatic beta-oxidation. Altogether, hepatic *Ass1* loss surprisingly does not prevent steatosis through cell-intrinsic lipid metabolism.

### Liver *Ass1* suppression increases whole body energy expenditure

Given known behavioral changes caused by hyperammonemia(*19*), we investigated whether these protective phenotypes could be attributed to changes in feeding or locomotion. GFP and mCre animals fed a HFD for 20 weeks were placed in metabolic cages for behavioral and metabolic assessment. Despite marked differences in body and fat mass (**Fig. S8A**), food intake and locomotor activity were unchanged (**Fig. S8B, C**). Modest, albeit variable, reduction in water intake was observed (**Fig. S8D**) but seemed unlikely to contribute to reduced body weight. Indirect calorimetry measurements, however, revealed increased energy expenditure during both daytime and nighttime in mCre animals compared to GFP controls (**Fig. S8E, F**), suggesting that hepatic *Ass1* loss increases basal metabolic rates. Respiratory exchange ratios were similar between groups (approximately 0.75), consistent with lipid as the primary fuel source (expected value of 0.7) on HFD (**Fig. S8G, H**)(*20*). Bomb calorimetry on feces collected from GFP and mCre animals revealed no significant changes in fecal output, fecal calories, or daily calories lost to feces (**Fig. S8I-K**). These results suggest that hepatic *Ass1* suppression increases basal whole body metabolic rates, primarily related to lipid metabolism.

### Loss of hepatic *Ass1* increases reliance on FAO in BAT

As livers in Cre animals exhibited no changes in FAO, we next measured FAO rates in other oxidative tissues known to use lipids for fuel, including BAT, skeletal muscle (tibialis anterior [TA] and soleus), and heart (*21–23*). We found that complete FAO rates were significantly increased in crude mitochondrial isolates from BAT of Cre animals compared to GFP controls (**Fig. 5A**), but not in skeletal muscle or heart (**Fig. S9A-C**). Consistent with increased activity, BAT from Cre animals exhibited smaller, more densely packed multilocular lipid droplets compared to GFP animals (**Fig. 5B**); however, this was not driven by changes in mitochondrial content or oxidative phosphorylation protein abundance (**Fig. S9D, E**). BAT primarily functions to generate heat through non-shivering thermogenesis, rapidly clearing plasma triglycerides through mitochondrial FAO upon activation(*24*). To test whether BAT FAO influenced phenotypes observed in Cre animals, we administered etomoxir (ETO, 15 mg/kg intraperitoneally [i.p.]), a carnitine palmitoyl transferase 1 (CPT1) inhibitor blocking the rate-limiting step of FAO, to HFD-fed GFP and Cre animals 10 days post-AAV (**Fig. 5C**). To our surprise, most Cre animals failed to survive a single ETO dose, while all GFP animals remained healthy (**Fig. 5C**), suggesting that FAO is essential upon acute hepatic *Ass1* loss. As ETO blocks FAO in both mitochondria and peroxisomes(*25*), we also treated mice with 10,12-tricosadiynoic acid (TDYA), an inhibitor of acyl-coA oxidase 1, to specifically inhibit peroxisomal FAO(*26*). TDYA (100 mg/kg daily oral gavage) did not affect HFD-fed GFP mice or further reduce survival of Cre animals (**Fig. 5C**), implicating reliance on mitochondrial FAO as the major reason for ETO-induced lethality. ETO sensitivity was greater at 12 days post AAV, with lethality occurring concomitantly with a rapid drop in core body temperature within 8 hours of a single ETO dose (**Fig. 5D, E**).

**Figure 5.**
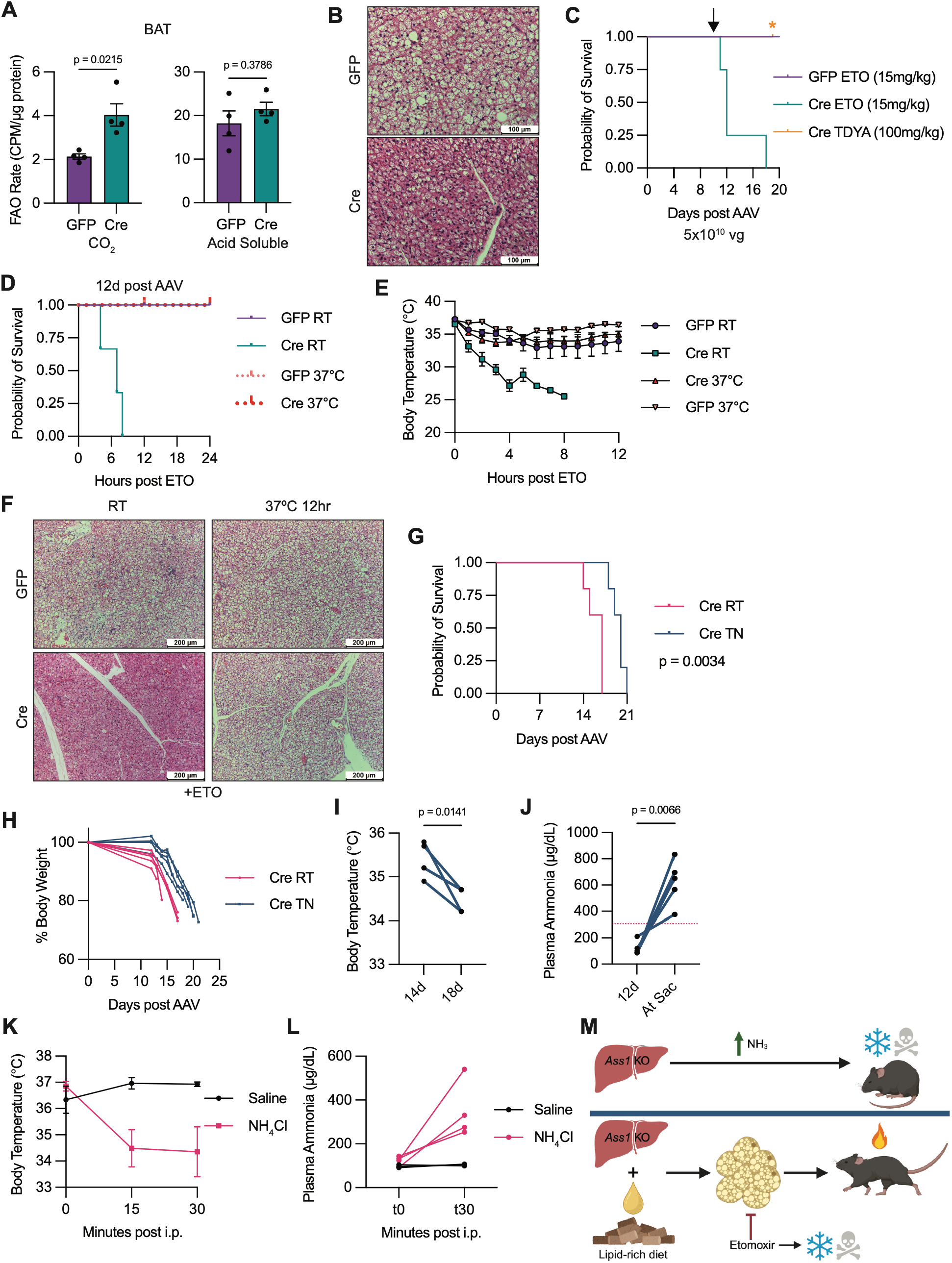
Hepatic *Ass1* loss activates BAT FAO for thermogenesis. **(A)** Mitochondria were isolated from BAT of GFP (n=4) and Cre (n=4) littermate pairs fed HFD for 14 days. Fatty acid oxidation rates were assessed by ^14^C-Palmitate oxidation to ^14^CO_2_ (complete oxidation) or ^14^C-acid soluble metabolites (incomplete oxidation). CPM, counts per minute. (**B**) Representative H&E images of BAT from GFP and Cre animals fed HFD for 14 days. (**C**) Survival curve of GFP and Cre animals maintained on HFD and treated with a single dose of etomoxir (ETO, 15mg/kg) on day 10 post AAV or daily treatment of TDYA (100mg/kg) beginning on day 10 post AAV. GFP ETO, n=4; Cre ETO, n=4; Cre TDYA, n=4. (**D-F**) GFP or Cre animals fed HFD were given a single dose of etomoxir (15mg/kg) on day 12 post AAV delivery then maintained at room temperature (RT) or on a 37°C heatpad. GFP RT, n=4; Cre RT, n=3; GFP 37°C, n=2; Cre 37°C, n=4. (**D**) Survival curve. (**E**) Body temperature. (**F**) Representative H&E images of BAT tissue. (**G-J**) Cre animals were fed SD and maintained at RT (n=5) or in a thermoneutral incubator at 30°C (TN, n=5) beginning at time of AAV delivery. (**G**) Survival curve. (**H**) Body weight change relative to body weight at AAV delivery. (**I**) Core body temperature of Cre TN animals measured by rectal thermometer at 14 and 18 days post AAV delivery. (**J**) Plasma Ammonia levels of Cre TN animals at 12 days post AAV delivery and at sacrifice. Dotted pink line represents average day 12 post AAV plasma ammonia levels from Cre animals housed at RT. (**K-L**) Wild-type animals were injected i.p. with saline (n=3) or NH_4_Cl (5mmol/kg, n=7). (**K**) Core body temperature. (**L**) Plasma ammonia levels measured before and 30 minutes post injection. Saline, n=3; NH_4_Cl, n=4. (**M**) Working model of *Ass1* regulation of whole-body energy homeostasis.

Given that thermogenesis is the primary function of BAT, we wondered whether external heat could support survival of Cre animals when FAO was inhibited. Indeed, providing access to a 37°C heat pad completely protected Cre animals from ETO-induced lethality by maintaining body temperature (**Fig. 5D, E**). Histological characterization of BAT from animals given ETO revealed markedly enlarged lipid droplets in heat rescued Cre animals, indicating reduced lipolysis in the absence of thermogenic demand (**Fig. 5F**). Together, these data indicate that rapid lethality observed upon hepatic *Ass1* loss may stem from an inability to maintain body temperature, as room temperature conditions constitute a mild cold stress for mice(*24*). Strikingly, thermoneutral housing (30°C), extended the lifespan of SD fed Cre animals by ∼30% (**Fig. 5G**). Thermoneutrality also preserved body weight and temperature for an extended period compared to room temperature conditions (**Fig. 5H, I**). Notably, plasma ammonia levels were lower in thermoneutral Cre animals than in room temperature-housed Cre animals at 12 days post-AAV delivery (**Fig. 1E**, **5J**); however, ammonia levels rose markedly by the time of sacrifice (**Fig. 5J**).

Infants with CTLN1 and other UCDs frequently present with low body temperature(*27–29*); therefore, we tested whether hyperammonemia itself could induce hypothermia. Consistent with a previous report(*30*), i.p. injection of wild-type mice with NH_4_Cl (5mmol/kg) caused an almost immediate drop in body temperature (**Fig. 5K**) concomitant with plasma ammonia levels of approximately 300 μg/dL, a level similar to Cre animals 12 days post AAV (**Fig. 5L**). Collectively, these data support a model in which *Ass1* loss-induced hyperammonemia triggers hypothermia, activating BAT thermogenesis and increasing the demand for dietary fat as fuel. This diverts lipids away from hepatic storage to confer protection against obesity, steatosis, and HCC (**Fig. 5M**).

## Discussion

Although the complete urea cycle is suppressed in liver disease and HCC (*2*, *6–8*), we show that ASS1 is distinct from other urea cycle enzymes(*12–14*) in that its loss protects against diet-induced obesity, liver steatosis, fibrosis, and HCC. This protection is driven by thermogenic BAT FAO activated in response to hyperammonemia-induced hypothermia. Given the diverse effects of high ammonia on the nervous system(*31*), we hypothesize that this hypothermic response is mediated by a neural mechanism(*32*). Consistent with this idea, plasma triglycerides were elevated in mCre animals (**Fig. 2I**), indicative of a systemic signal promoting fat mobilization(*33*). Critically, this thermogenic demand necessitates dietary fat for survival, as inadequate lipid supply was fatal. We further show that targeting hepatic *Ass1* reverses obesity even after disease onset. Careful titration of hepatic *ASS1* targeting, perhaps through liver-targeted nanoparticles or AAV vectors(*34*), presents a potential therapeutic strategy, although the relatively low BAT abundance in human adults compared to rodents warrants careful consideration (*35*).

Chronically activated BAT likely underlies the observed hepatic lipid phenotypes. BAT rapidly clears circulating triglycerides through mitochondrial FAO(*36*), and we propose this creates a sustained systemic demand for lipid fuel that reduces fatty acid availability for hepatic storage. Since steatosis drives liver disease progression(*37*, *38*), diversion of lipids towards BAT thermogenesis may render the liver less permissive to disease initiation, explaining protection from liver disease and HCC observed in mCre animals despite continued high fat feeding.

The reciprocal relationship between ammonia and body temperature remains incompletely understood. A recent report implicates the transient receptor potential (TRP) ion channel family in modulating the magnitude of ammonia’s hypothermic effect(*39*), yet the precise mechanism by which hyperammonemia regulates body temperature remains unknown. Intriguingly, our observation that thermoneutrality reduces plasma ammonia levels (**Fig. 5J**) appears to contradict previous reports proposing therapeutic hypothermia as a treatment for hyperammonemia(*40*). Additional work is required to resolve this apparent discrepancy.

Although *ASS1* loss promotes growth in established cancer cell lines(*4*, *5*, *41*), our data support a model where systemic metabolic reprogramming instead protects against tumor initiation(*42*). Notably, we occasionally observed extremely large tumors in mCre livers (**Fig. 3N, P**), suggesting that in fully transformed tumor cells, *Ass1* loss may instead promote tumor growth, consistent with prior reports.

These findings also have important clinical implications for CTLN1 patients. Although liver transplant and gene therapy are curative(*43*, *44*), outcomes improve in older infants who are given time to gain weight(*45*). We demonstrate that high dietary fat significantly extends the lifespan of Cre mice (**Fig. 1C**) by providing fuel for hyperactivated BAT thermogenesis. Given that BAT constitutes ∼5% of human infant mass(*46*), optimizing fat intake may expand the therapeutic window for severe CTLN1. For patients managed with ammonia scavengers and low-protein diet, current guidelines require lifelong liver surveillance(*11*); however, our data suggest CTLN1 patients might face reduced risk for liver disease and HCC compared to other UCDs like ASL and citrin deficiency(*12*, *13*). Consistent with this, liver biopsies and postmortem samples revealed relatively normal liver histology in CTLN1 patients compared to CPS1- or ASL-deficient patients(*47*). These findings support larger-scale studies of hepatic manifestations across individual UCDs that may inform updated surveillance guidelines.

## Acknowledgements

The authors thank the entire Simon laboratory for comments and discussions on the manuscript. We thank Michelle Burrows and the Lazar laboratory for assistance with thermoneutral experiments. BioRender was used for the creation of graphical and schematic models. AI-assisted tools were used for grammar and style review. We thank the Rodent Metabolic Phenotyping Core (RRID: SCR_022427), supported in part by the National Institutes of Health grant S10-OD025098, the Cox Institute, and the Institute for Diabetes, Obesity, and Metabolism (Perelman School of Medicine, University of Pennsylvania) for performing indirect calorimetry and fecal bomb calorimetry experiments. We thank the Molecular Pathology and Imaging Core (RRID: SCR_022420) supported by the Center for Molecular Studies in Digestive and Liver Diseases (P30DK050306) for assistance with histology. We thank the Molecular Profiling Facility (RRID: SCR_024999) for assistance with RNA sequencing analysis.

## Funding

Ludwig Cancer Research Institute (MCS) NCI R35CA220483 (MCS)

National Cancer Institute (NCI) 5T32CA009140 (LCK and NPL)

American Cancer Society Postdoctoral Fellowship PF-23-1034739-01-TBE (LCK)

Damon Runyon Postdoctoral Fellowship DRG2497 (NPL)

NCI F30 CA271654 (NJC)

NIEHS P30 ES013508 (CM and JPX)

## Author Contributions

Conceptualization: LCK, MCS

Formal Analysis: LCK

Funding Acquisition: MCS

Investigation: LCK, NPL, AZ, XC, XH, JPX, NJC, DZ

Methodology: LCK, NPL, JPX, CM

Resources: CM, ZA

Supervision: ZA, MCS

Validation: LCK, AZ

Visualization: LCK

Writing-original draft: LCK

Writing-review & editing: LCK, BK, MCS

## Data and materials availability

RNA-sequencing data will be deposited to GEO upon publication. Lipidomics and metabolomics data will be deposited on Metabolomics Workbench upon publication. All other materials generated in this study will be available upon request.

