## supplement for "Hepatic urea cycle suppression drives brown fat thermogenesis to counter obesity and liver disease"

**The PDF file includes:**

Materials and Methods
Figs. S1 to S9
Tables S1 and S2

### **Materials and Methods**

Mouse experiments

All animal experiments were reviewed and approved by the Institutional Animal Care and Use Committee (IACUC) at the University of Pennsylvania. C57BL/6 *Ass1*$^{flox/flox}$ animals were a generous gift from Dr. Eleonore S. Köhler(*48*). Unless indicated, all experiments were performed on male mice. AAV8-TBG-GFP (Addgene, #05535) or AAV8-TBG-Cre (Addgene, 107787) were delivered by retroorbital injection at indicated ages. For obesity and liver steatosis experiments, animals were fed a HFD with 60% kcal from fat (TD. 06414). For MASH development, animals were fed the Gubra-Amylin NASH diet (Research Diets, D09100310). For HCC experiments, 14-day old animals were administered DEN (25 mg/kg) intraperitoneally (i.p.), then AAV and HFD were started at 6 weeks old. Etomoxir (15 mg/kg, MCE HY-50202A) was diluted in saline and delivered i.p. at indicated time points. 10,12-tricosadiynoic acid (100 mg/kg, Millipore Sigma, 91445) was diluted in olive oil and delivered daily by oral gavage. $NH_4Cl$ (5mmol/kg) was diluted in saline and delivered i.p. to untreated *Ass1*$^{flox/flox}$ animals. Thermoneutrality experiments were performed in climate-controlled rodent incubators (Powers Scientific) maintained at 30°C, with interior temperature monitoring by digital thermometers. Internal core body temperatures were recorded using a digital thermometer (Oakton Instruments, Temp 10T Thermocouple) and rectal thermocouple probe (Physitemp, RET-3 probe). For short term warming experiments, cages were set on top of a far-red infrared warming pad (Kent Scientific #RT-JR and #RT-0502) set to 37°C. Animals were sacrificed if core body temperature dropped 10 degrees or body weight dropped more than 20%.

Metabolic parameters, energy expenditure, and locomotor activity were analyzed using the Promethion indirect calorimetric system (Sable Systems). Measurements were collected for the dark (from 7 p.m. to 7 a.m.) and light (from 7 a.m. to 7 p.m.) periods under *ad libitum* feeding. Mice were allowed to acclimatize to the individual metabolic cages for 48 hours before measurements were recorded. Analysis was performed with CalR2 web-based tool(*49*). Body composition was analyzed by DEXA PIXImus mouse densitometer. Fecal bomb calorimetry was performed on feces collected over a 24-hour period from singly housed animals on the IKA C3000 Calorimeter.

Histology, Immunohistochemistry, and Immunofluorescence

Mouse tissues were fixed in 4% PFA, or embedded in OCT right after collection, and submitted to the Molecular Pathology & Imaging Core (MPIC) at the University of Pennsylvania for processing, sectioning, and hematoxylin & eosin (H&E) staining. For IF, slides were deparaffinized, rehydrated, quenched in 0.6% hydrogen peroxide/methanol for 15 min. Citrate (10 mM, pH 6.0) buffer was used for antigen retrieval. Slides were incubated with primary antibodies overnight at 4°C, and then with AlexaFluor conjugated secondary antibodies the next day. Incubations with antibodies directly conjugated to fluorophores were performed sequentially. Slides were mounted with VECTASHIELD Antifade Mounting Media with DAPI.

Oil Red O Staining

Frozen sections were fixed in 4% paraformaldehyde (Thermo Fisher Scientific, 28908) for 15 min. Samples were stained with Oil Red O (Sigma-Aldrich, P6148) for 30 min at room temperature. Oil Red O working solution was prepared by diluting 3.5 mg/mL stock (in 100% isopropanol) 6:4 with distilled water. The slides were rinsed in distilled water and counterstained with hematoxylin before mounting with Vectashield and imaged using a Leica DM2000 bright-field microscope.

Sirius Red Staining

Paraffin-embedded tissue slides were deparaffinized, rehydrated and incubated with pre-warmed Bouin's solution at 55 °C for 1 h. The slides were then washed and incubated in 0.1% fast green (Sigma) solution for 10 min, then in 1% acetic acid for 2 min. Slides were then stained in 0.1% Sirius Red (Sigma) solution for 30 min, dehydrated and then mounted for examination.

RNA isolation, RT and qRT-PCR

Total RNA was isolated using RNeasy Mini Kit (Qiagen). cDNA was synthesized using a High-Capacity RNA-to-cDNA Master Mix (Applied Biosystems). qRT-PCR was performed using TaqMan Universal PCR Master Mix (Thermo Fisher) or Power SYBR Green Master Mix (Applied Biosystems) on a QuantStudio 6 (Applied Biosystems). Expression levels were normalized to*18S* rRNA or Actin. Primer sequences and catalog numbers are provided in Table S2.

RNA-sequencing and data analysis

Total RNA was extracted from snap frozen tissue samples using RNeasy Mini Kit (Qiagen). RNA quality test, library construction and sequencing were performed by Novogene Corporation. Data was analyzed at the Molecular Profiling Facility at the University of Pennsylvania. Briefly, Fastq files were checked for quality using FastQC and qualimap. Alignment was performed using the STAR aligner under default settings with the mm10 reference genome. Raw counts of gene transcripts were obtained from the resulting bam files using feature Counts. The raw count matrix was subsequently imported into R-studio (R version 3.3.3) and used as input for DESeq2 following the vignette of the package for normalization and differential gene expression analysis. Salmon/Sailfish was used in parallel to normalize and quantitate gene expression in transcripts per million (TPM) through quasi-alignment. GSEA (http://software.broadinstitute.org/gsea/index.jsp) was run for the contrast in pre-ranked mode using the DESeq2 statistic as the ranking metric. Mouse gene symbols were mapped to human gene orthologs using Ensembl's BioMart (http://www.ensembl.org/biomart/martview/).

Plasma Collection and Tests

Blood was collected by retroorbital bleeding in heparinized tubes. Samples were centrifuged at 10,000xg for 5 min and plasma was collected. Plasma triglycerides were measured using the Infinity Triglyceride Reagent. Plasma Ammonia was measured using the Infinity Ammonia Reagent or by IDEXX Chemistry Analyzer. Absorbance was read on a Tecan plate reader. ALT levels were measured using the ALT Activity Kit (Invitrogen #EEA001) or IDEXX Chemistry Analyzer. AST levels were measured using the AST Activity Assay Kit (Abcam #ab105135).

VLDL-TG Secretion

Animals were fasted for 4 hours, and blood was sampled to determine baseline plasma triglyceride levels. Then animals were injected with Poloxamer 407 (670 mg/kg, MedChem Express, HY-D1005) to inhibit lipoprotein lipase activity. Blood samples were collected via the tail vein at 1, 2, and 4 hours post drug administration. Plasma was collected and analyzed for triglyceride abundance using the Infinity Triglyceride reagent.

Lipidomics

Lipids were extracted as recently described*(50)*. Flash-frozen liver tissue was pulverized under liquid nitrogen, and approximately 10 mg of tissue powder was aliquoted into Eppendorf tubes. To each tube, 1 mL of cold (−80°C) 80% methanol (MeOH) and 20 µL of SPLASH LIPIDOMIX internal standard mixture (Avanti Polar Lipids, 330707; diluted 1:1 (v/v) in MeOH) were added. Samples were pulse sonicate to disrupt the membranes, vortexed and transferred to 10 mL glass Pyrex tubes. The Eppendorf tubes were rinsed with 200 µL MeOH, which was combined with the corresponding sample. Optima-grade water (1.2 mL) was added, followed by vortexing. Formic acid (100 µL, 0.1%) and 5 mL methyl *tert*-butyl ether (MTBE) were then added. Tubes were capped and shaken for 20 min on a mechanical shaker (high setting). Phase separation was induced by centrifugation at 600 × g for 20 min using low acceleration and deceleration settings to prevent glass breakage. The upper organic phase (MTBE/MeOH), containing lipids, was collected into new 10 mL glass Pyrex tubes and evaporated to dryness under nitrogen. Dried extracts were reconstituted in 200 µL MTBE/MeOH (1:3, v/v). Extraction blanks were prepared in the same way using solvents only, without tissue. A pooled quality control (QC) sample was prepared by combining 50 µL from each reconstituted sample. All samples were processed in a single batch. An aliquot (100 µL) of each sample was transferred to amber glass HPLC vials with deactivated inserts for analysis.

Lipids were analyzed by liquid chromatography high-resolution mass spectrometry (LC-HRMS) as recently described*(50)*. Lipids were separated on an Accucore C18 column (2.1 × 100 mm, 2.6 µm; Thermo Fisher Scientific, #17126-102130) maintained at 35 °C on a Dionex UltiMate 3000 quaternary UHPLC system equipped with a refrigerated autosampler (10°C). Solvent A was acetonitrile/water (1:1, v/v) and solvent B was acetonitrile/isopropanol/water (10:88:2, v/v/v), both containing 10 mM ammonium formate and 0.1% formic acid. The flow rate was 0.4 mL/min, and the gradient was: 0–1 min, 90% A; 4 min, 60% A; 12 min, 25% A; 21–24 min, 1% A; 24.1–28 min, 90% A.

Samples (2 µL injection) were analyzed on a Q Exactive HF mass spectrometer (Thermo Fisher Scientific) in positive ion mode and separately in negative ion mode, and the column effluent was directed to the mass spectrometer from 2 to 23 min and to waste for the remainder of the run. Data were acquired in full scan/data-dependent MS/MS (Top 10) mode. Full scan settings were: resolution 120,000, AGC target $1 \times 10^6$, maximum injection time 120 ms, and scan range *m/z* 250–1800. MS/MS spectra were acquired at resolution 15,000 with AGC target $1 \times 10^5$, maximum injection time 25 ms, and stepped normalized collision energy (NCE) of 25, 30, and 35. Source parameters were: spray voltage 4000 V (positive) and 3500 V (negative), capillary temperature

285 °C, probe heater temperature 375 °C, sheath gas 45 (positive)/55 (negative), auxiliary gas 15 (positive)/20 (negative), sweep gas 1, and S-lens RF level 45. Features detected in the extraction blanks above an intensity of $1 \times 10^5$ were placed on an exclusion list. External mass calibration was performed approximately every 5 days with the Thermo calibration mixture. Samples were analyzed in random order, with the pooled QC injected every 10 samples.

Lipids were identified and area under the peak integrated with LipidSearch 4.2 (Thermo Fisher Scientific) using product search, with a precursor tolerance of 5 ppm, a product ion tolerance of 10 ppm, a retention time tolerance of 0.15 min, and a signal-to-noise threshold of 5. Lipid species were retained when their coefficient of variation in the pooled QC samples was below 30% and their QC/blank intensity ratio was above 10. Acyl chains separated by an underscore indicate that their *sn*-positions were not determined. The deuterated internal standards from SPLASH LIPIDOMIX were identified in LipidSearch, and the peak area of each lipid was normalized to the internal standard of the same class (PC, PE, PG, PI, PS, LPC, LPE, DG, TG, and SM) and to tissue weight; amounts are reported as pmol/mg tissue. Because one internal standard was used per class, differences in response between molecular species of the same class were not corrected, and these amounts are semi-quantitative. Ceramides and acylcarnitines, which have no matched standard in SPLASH LIPIDOMIX, were normalized to the closes deuterated standard as retention time, LPC and SM, and are reported as relative abundance. Total class abundances were obtained by summing all species in each class. The 18:1/18:0 desaturation index was calculated as summed abundance of triglyceride species containing 18:1 divided by that of species containing 18:0.

Metabolite Profiling of Plasma and Liver

Metabolism of plasma or flash frozen liver was stopped by addition of cold 40% acetonitrile/40% methanol/20% water. Tissues were lysed via three freeze/thaw cycles, centrifuged to remove protein (15 min, 4 °C, 15000 x g), and metabolite supernatant taken for analysis. Data acquisition was performed by hydrophilic interaction chromatography on a Vanquish Flex UHPLC (Thermo) system coupled to a Q Exactive mass spectrometer (Thermo). The mass spectrometer (MS) was operated in positive or negative ion mode. Analytes were separated on a Sequant ZIC-pHILIC HPLC column (5 µm x 2.1 mm x 150 mm) at 25 °C. Mobile phase A composition was 10 mM ammonium acetate pH 9 and mobile phase B was 100% ACN. The LC gradient was: 0 min: 90% B; 20 min: 40% B; 20-20.5 min: 90% B; 20.5-28 min: 90% B. The flow rate was 150 µl/min with a sample injection volume of 2 µl. The MS was operated in full MS with the following parameters: resolution (140,000), AGC target (1e6), maximum IT (100 ms), and scan range (55 – 825 m/z). Data dependent MS/MS (dd-MS2) used the following parameters: resolution (17,500), AGC target (1e5), maximum IT (50 ms), loop count (5), topN (5), isolation window (1.5 m/z), stepped NCE (30, 60, 90), minimum AGC (8e3), and dynamic exclusion (4s). Data was collected by the Thermo Fisher software Xcalibur (v4.5.474.0). Peak picking as performed in El-MAVEN for positive and negative ion mode, and clustering performed using Metaboanalyst (https://metaboanalyst.ca/).

Free Fatty Acids and Glycerol Measurements

Flash frozen liver tissue was pulverized in liquid nitrogen and homogenized in PBS. Free fatty acids were measured with the Free Fatty Acid Assay Kit (Sigma MAK466) and glycerol was measured with the Glycerol Assay Kit (APEXbio #K2134).

### Hepatocyte Isolation

Hepatocytes were isolated according to modified previously published protocols(*51*) . Briefly, the inferior vena cava was exposed in anesthetized mice to insert a catheter needle delivering liver perfusion buffer (HBSS no $Ca^{2+}$, no $Mg^{2+}$, no phenol red; 0.5mM EDTA; 25mM HEPES) via a peristaltic pump. Upon proper perfusion, 10 mL of warmed collagenase IV (0.1mg/mL) in HBSS with 25 mM HEPES was delivered at a rate of 1 mL/min. When perfusion was finished, the liver was removed and ruptured to release hepatocytes. Cells were filtered through a 70um filter two times, then centrifuged (200xg) through a 45% Percoll gradient for 10 min at 4°C. Live hepatocytes in the pellet were resuspended in DMEM low glucose with 5% FBS and 1% Penicillin-Streptomycin.

### Fatty Acid Oxidation Rate Measurement

Fatty acid oxidation rates were measured according to previously published protocols with slight modifications(*52*). Cells or crude mitochondrial isolates were incubated in 400µL reactions with 0.2mM 0.5µCi/ml [1-$^{14}$C]-palmitic acid (Revvity NEC075H050UC) for 30 min at 37°C. Reactions were stopped with 200µL of 3M Perchloric acid and $CO_2$ was trapped with 20µL of 1M NaOH on a Whatman paper cap for 1h at room temperature. The filter paper was removed and acidified medium was centrifuged (10,000 xg for 10min). The radioactivity of the trapped $CO_2$, which corresponds with complete oxidation, and the radioactivity of the acid soluble metabolites contained in the supernatant, which is equivalent to incomplete oxidation, were measured in a scintillation counter.

### Statistics

Statistical analyses were performed with GraphPad Prism using a two-tailed Student's *t*-test (unpaired unless otherwise noted) to compare two groups, one-way ANOVA with Tukey's post hoc test to compare more than two group, and Logrank (Mantel-Cox) test to compare survival curves. The data are shown as the mean ± s.e.m. unless stated otherwise. All data points represent individual biological samples. Lipidomics and metabolomics data were $\log_2$-transformed before univariate analysis. P-values for volcano plots were calculated using unpaired two-tailed Student's t-test. Heatmaps show row z-scores with hierarchical clustering if indicated.

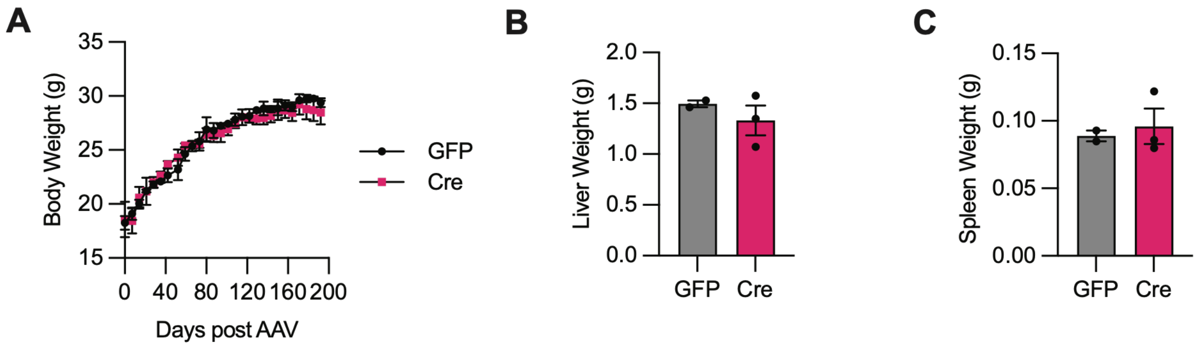


**Fig. S1. Mice with heterozygous deletion of hepatic *Ass1* are phenotypically unremarkable.** (**A-C**) AAV8-TBG-GFP control (n=2) or -Cre ($5x10^{10}$ viral genomes, vg/mouse; n=3) was administered to *Ass1*$^{fl/+}$ mice at 6 weeks of age and animals were sacrificed after 24 weeks. (**A**) Body weight over time post AAV delivery. (**B**) Liver weight at sacrifice. (**C**) Spleen weight at sacrifice.

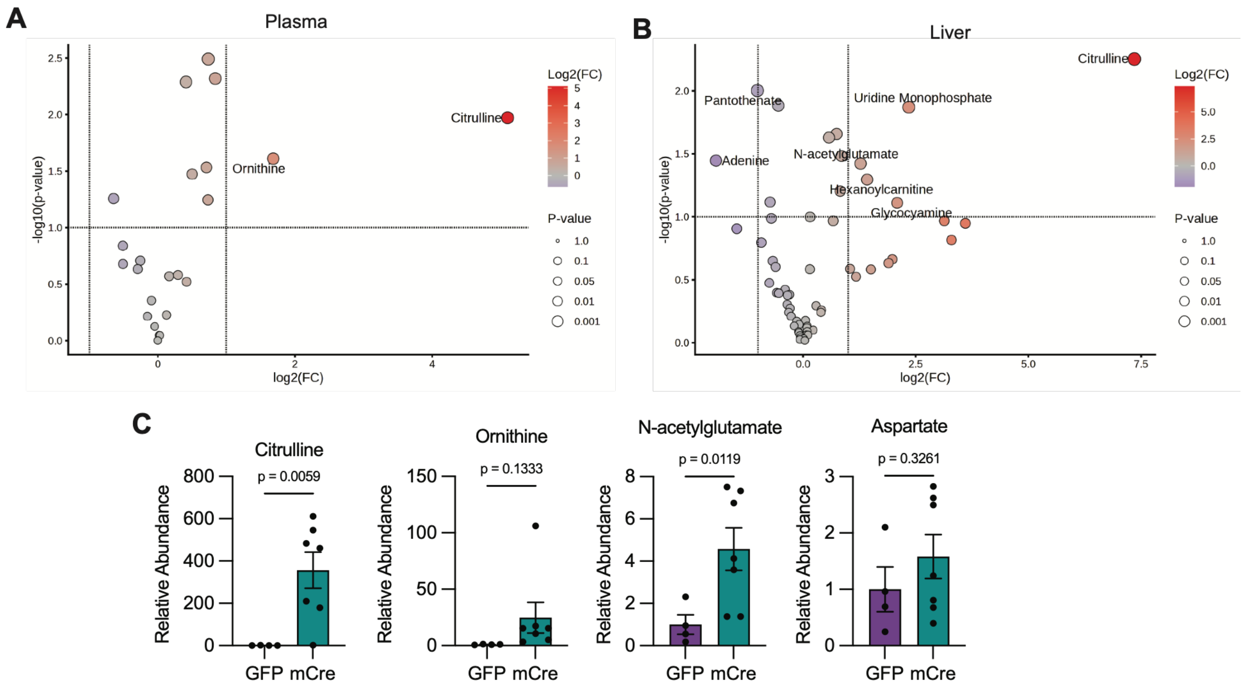


**Fig. S2. Metabolomic analysis of mCre plasma and liver.**

(**A**) Volcano plot of metabolites detected by LC/MS in plasma of mCre/GFP mice after 20 weeks on HFD. Significantly altered metabolites are labeled. GFP, n=4; mCre, n=7. (**B**) Volcano plot of metabolites detected by LC/MS in livers of mCre/GFP mice after 20 weeks on HFD. Significantly altered metabolites are labeled. (**C**) Relative abundance of urea cycle related metabolites in livers of GFP and mCre mice. Fold change, FC.

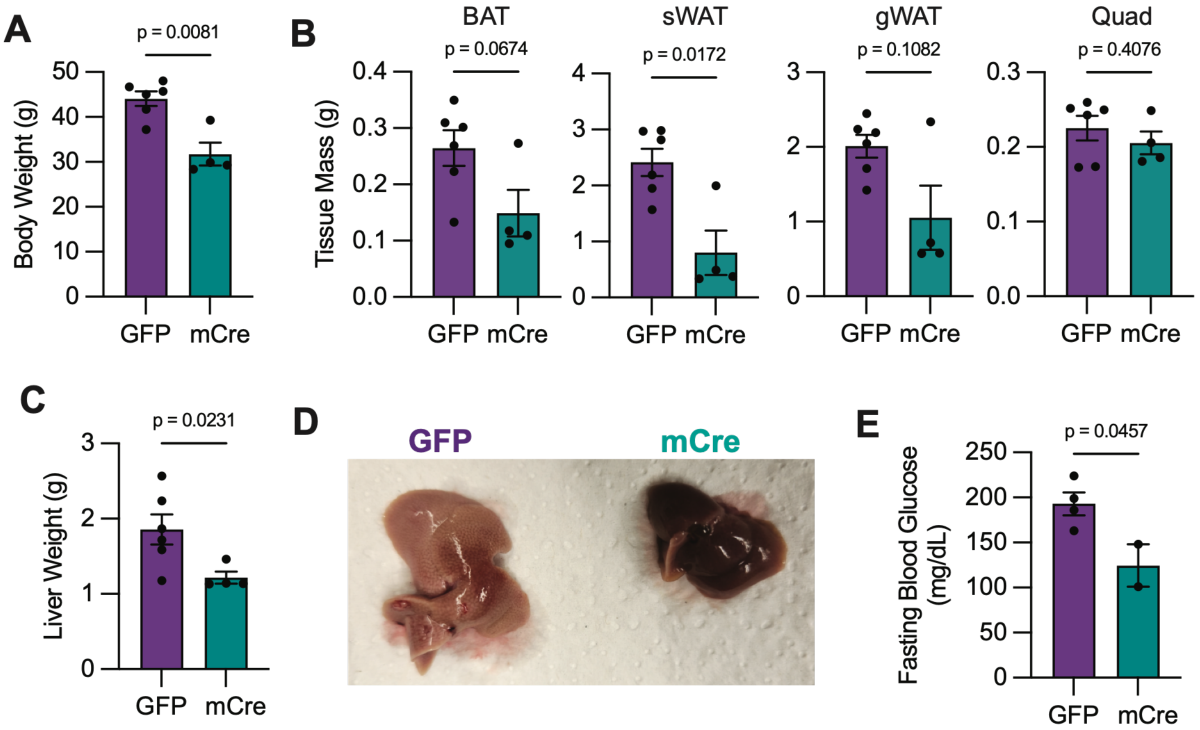


**Fig. S3. mCre animals are protected from obesity and severe hepatosteatosis after 36 weeks on HFD.**

(**A-E**) AAV8-TBG-GFP control (n=6) or -Cre (mCre, 2.5x$10^{10}$ vg/mouse; n=4) was administered to *Ass1*$^{fl/fl}$ mice at 6 weeks of age. HFD feeding was started at the time of AAV delivery. Animals were monitored for 36 weeks, then sacrificed for analysis. (**A**) Body weight at sacrifice. (**B**) Tissue weights. BAT, brown adipose tissue; gWAT, gonadal white adipose tissue; sWAT, subcutaneous white adipose tissue; Quad, quadriceps. (**C**) Liver weight. (**D**) Representative images of gross livers. (**E**) Fasting blood glucose. GFP, n=4; mCre, n=2.

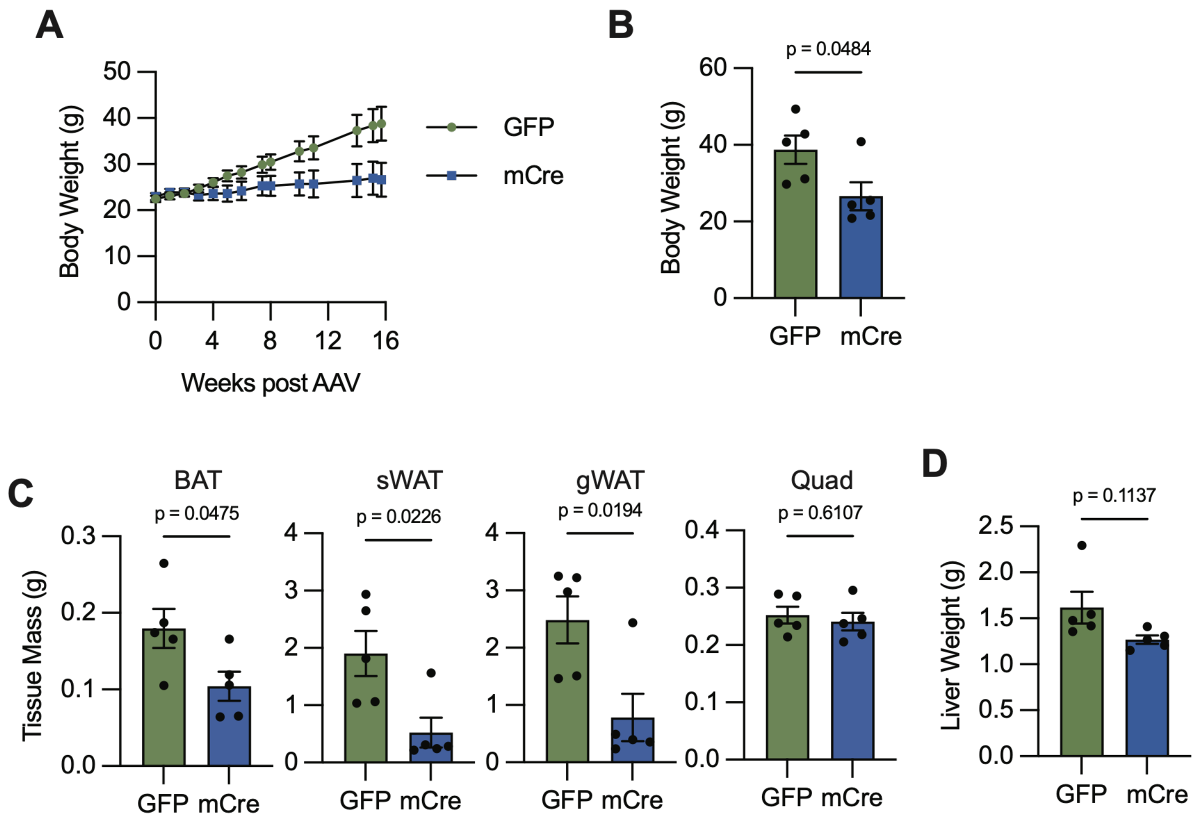


**Fig. S4. Mosaic *Ass1* loss in liver prevents obesity induced by ketogenic diet.**
(**A-D**) AAV8-TBG-GFP control (n=5) or -Cre (mCre, $2.5x10^{10}$ vg/mouse; n=5) was administered to *Ass1*$^{fl/fl}$ mice at 6 weeks of age. Ketogenic diet was started at the time of AAV delivery. Animals were monitored for 15 weeks, then sacrificed for analysis. (**A**) Body weight on ketogenic diet. (**B**) Body weight at sacrifice. (**C**) Tissue weights. (**D**) Liver weight.

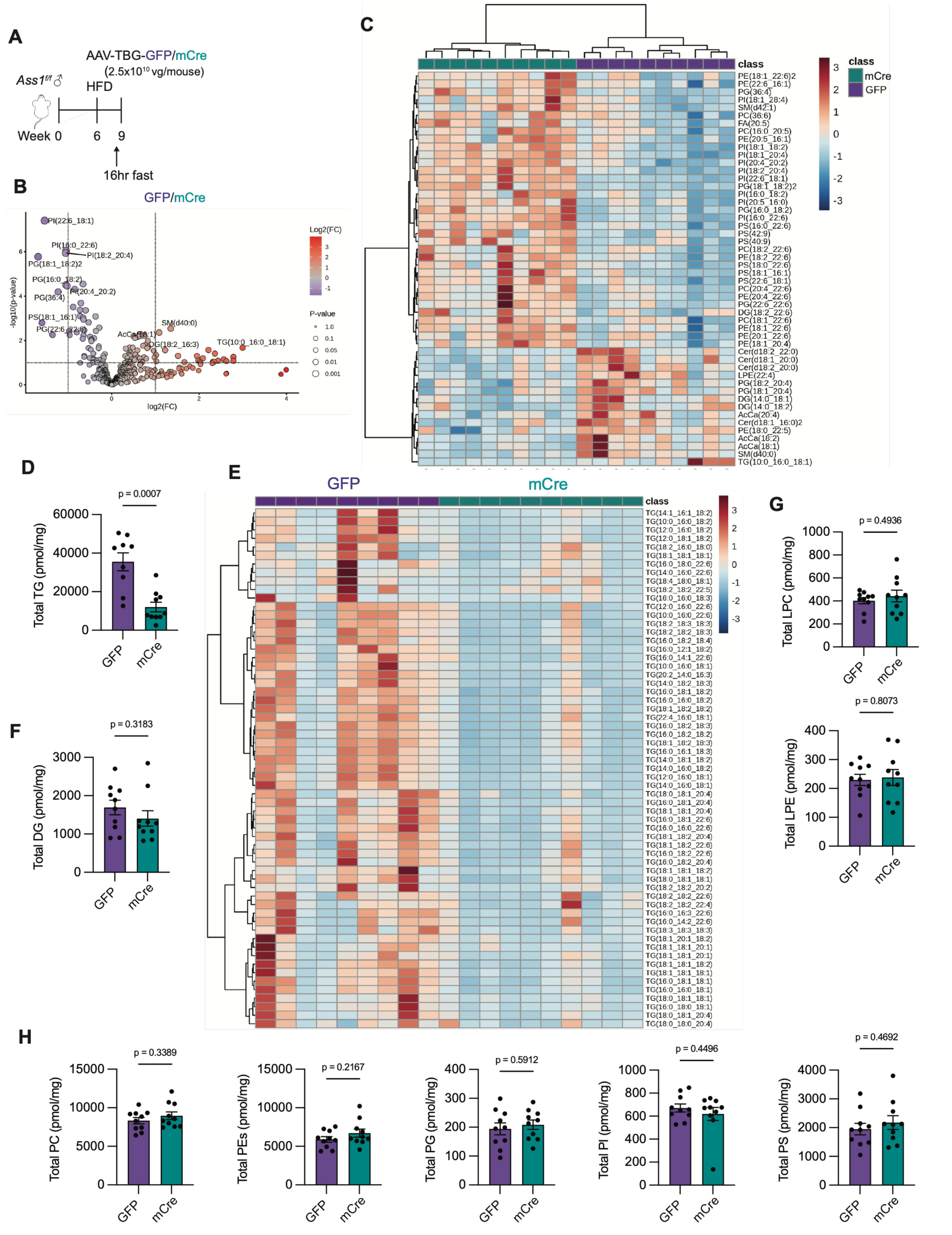


**Fig. S5. *Ass1* supports hepatic triglyceride stores.**

(**A-J**) AAV8-TBG-GFP control or -Cre (mCre, 2.5x$10^{10}$ vg/mouse) was administered to *Ass1$^{fl/fl}$* mice at 6 weeks of age. HFD was started at the time of AAV delivery. Animals were sacrificed after 3 weeks and livers were subjected to lipidomics analysis. N = 10 per condition. (**A**) Experimental schema. (**B**) Volcano plot of all lipid species detected by lipidomics. (**C**) Top 50 most significantly altered lipid species with clustering analysis. (**D**) Total abundance of all triglyceride species. (**E**) Heatmap of individual triglycerides detected by lipidomics. (**F**) Total abundance of all diglyceride species. (**G**) Total abundance of lysophospholipids including lysophosphatidylcholines, LPC, and lysophosphatidylethanolamines, LPE. (**H**) Total abundance of phospholipids including phosphatidylcholines, PC, phosphotidylethanolamines, PE, phosphatidylglycerols, PG, phosphatidylinositols, PI, and phosphatidylserines, PS.

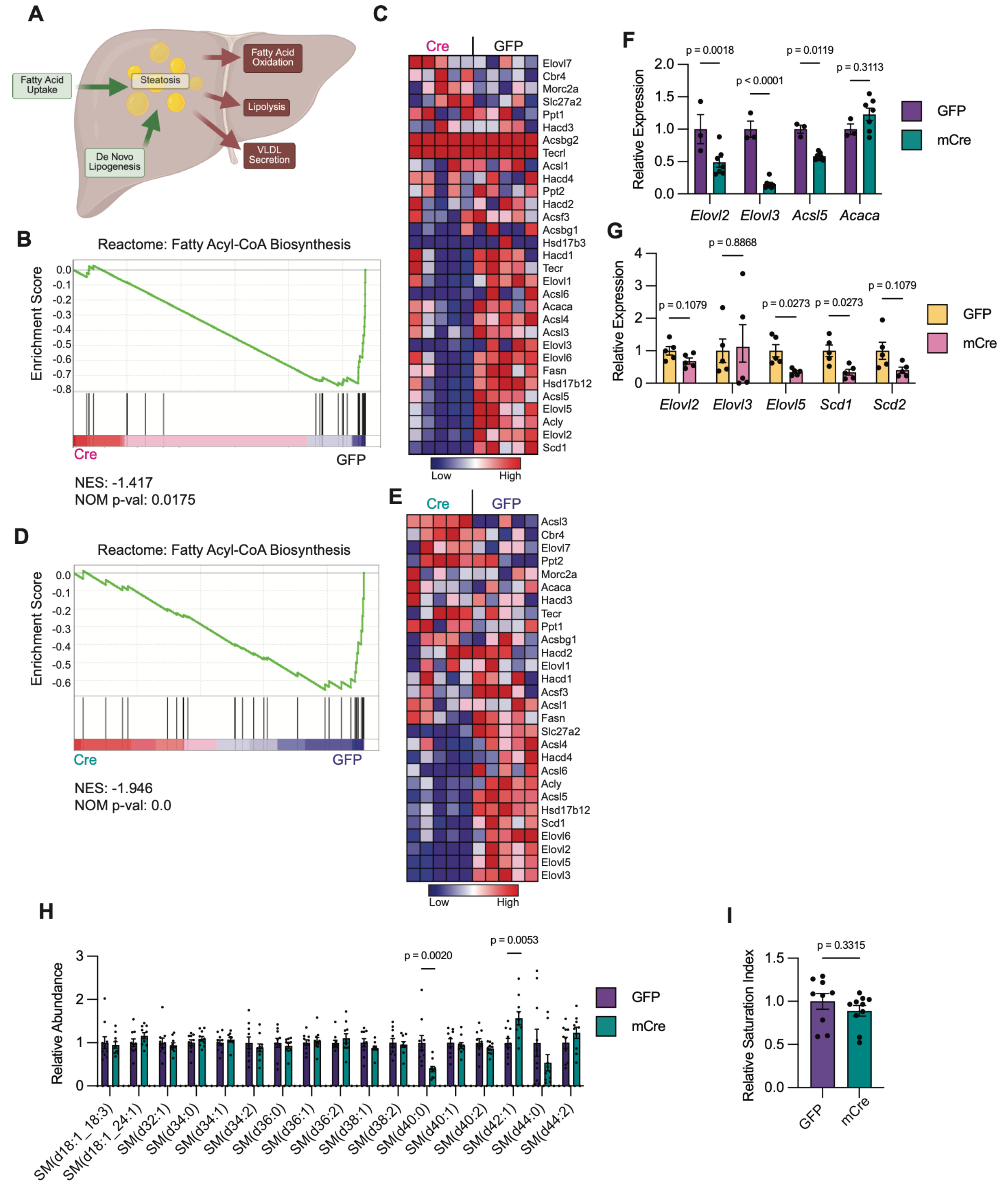


**Fig. S6. Fatty acyl-CoA biosynthesis genes are suppressed by loss of *Ass1* in liver.**
(**A**) Schematic depicting lipid metabolism pathways related to liver steatosis. (**B-C**) Gene set enrichment analysis for the Reactome Fatty Acyl-CoA Biosynthesis pathway in livers of GFP (n=5) and Cre (n=5) animals fed SD 13 days post AAV delivery. (**B**) Enrichment plot. (**C**) Heat map of all genes in gene set. (**D-E**) Gene set enrichment analysis for the Reactome Fatty Acyl-

CoA Biosynthesis pathway in livers of GFP (n=5) and Cre (n =5) animals fed HFD 3 weeks post AAV delivery. (**D**) Enrichment plot. (**E**) Heat map of all genes in gene set. (**F**) qRT-PCR analysis of livers from GFP (n=3) and mCre (n=7) animals on HFD for 20 weeks. (**G**) qRT-PCR analysis of livers from GFP (n=5) and mCre (n=5) animals on GAN diet for 25 weeks. (**H**) Relative abundance of individual sphingomyelin, SM, species from livers described in Fig. S5. (**I**) Ratio of the total abundance of 18:1 acyl chains relative to 18:0 acyl chains detected in triglyceride species from livers described in Fig. S5.

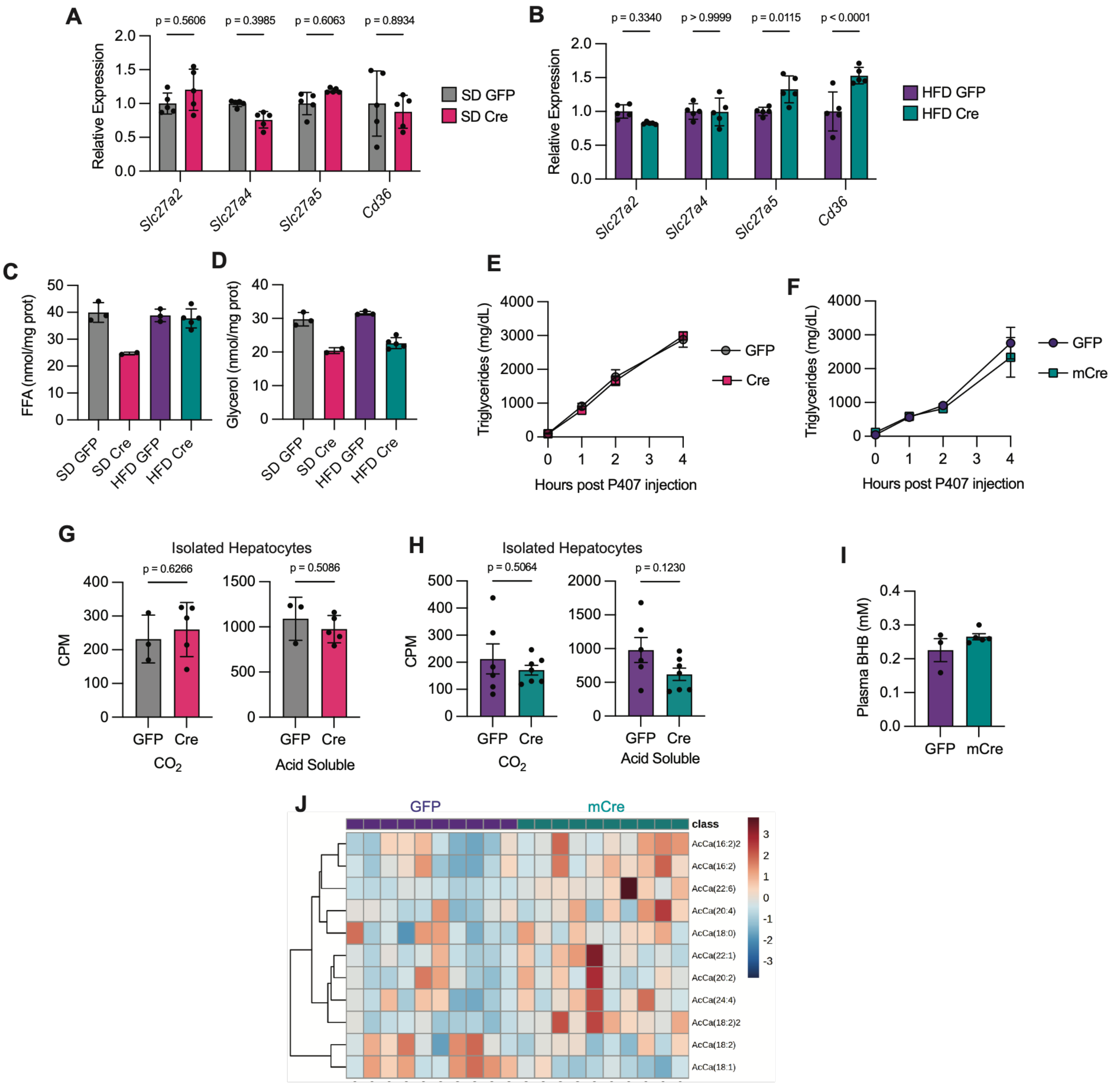


**Fig. S7. Protection from steatosis is independent of liver lipid metabolism.**

(**A**) Relative mRNA expression of fatty acid transporters as assessed by RNA sequencing in livers of GFP (n=5) or Cre (n=5) animals on SD collected 13 days post AAV delivery. (**B**) Relative mRNA expression of fatty acid transporters as assessed by RNA sequencing in livers of GFP (n=5) or Cre (n=5) animals on HFD collected 3 weeks post AAV delivery. (**C-D**) Livers from GFP or Cre animals fed either SD or HFD for 13 days post AAV delivery were analyzed for free fatty acids, FFA (**C**), or glycerol (**D**). SD GFP, n=3; SD Cre, n=2; HFD GFP, n=3; HFD Cre, n=5. (**E-F**) VLDL-triglyceride secretion rates in GFP (n=6) and Cre (n=6) animals fed SD for 13 days (**E**) or GFP (n=4) and mCre (n=3) animals fed HFD for 5 weeks (**F**). (**G-H**) Fatty acid oxidation rates assessed by $^{14}$C-Palmitate oxidation to $^{14}CO_2$ (complete oxidation) or $^{14}$C-acid soluble metabolites (incomplete oxidation) in isolated hepatocytes from GFP and Cre animals fed SD (GFP, n=3; Cre, n=5; **G**) or HFD (GFP, n=6; Cre, n=7; **H**) for 13 days. CPM, counts per minute. (**I**) Plasma betahydroxybutyrate, BHB, levels in GFP (n=3) and mCre (n=5) animals after 5 weeks on HFD measured by LC/MS**.** (**J**) Liver acyl-carnitine levels measured by LC/MS in GFP and mCre animals described in Fig. S5.

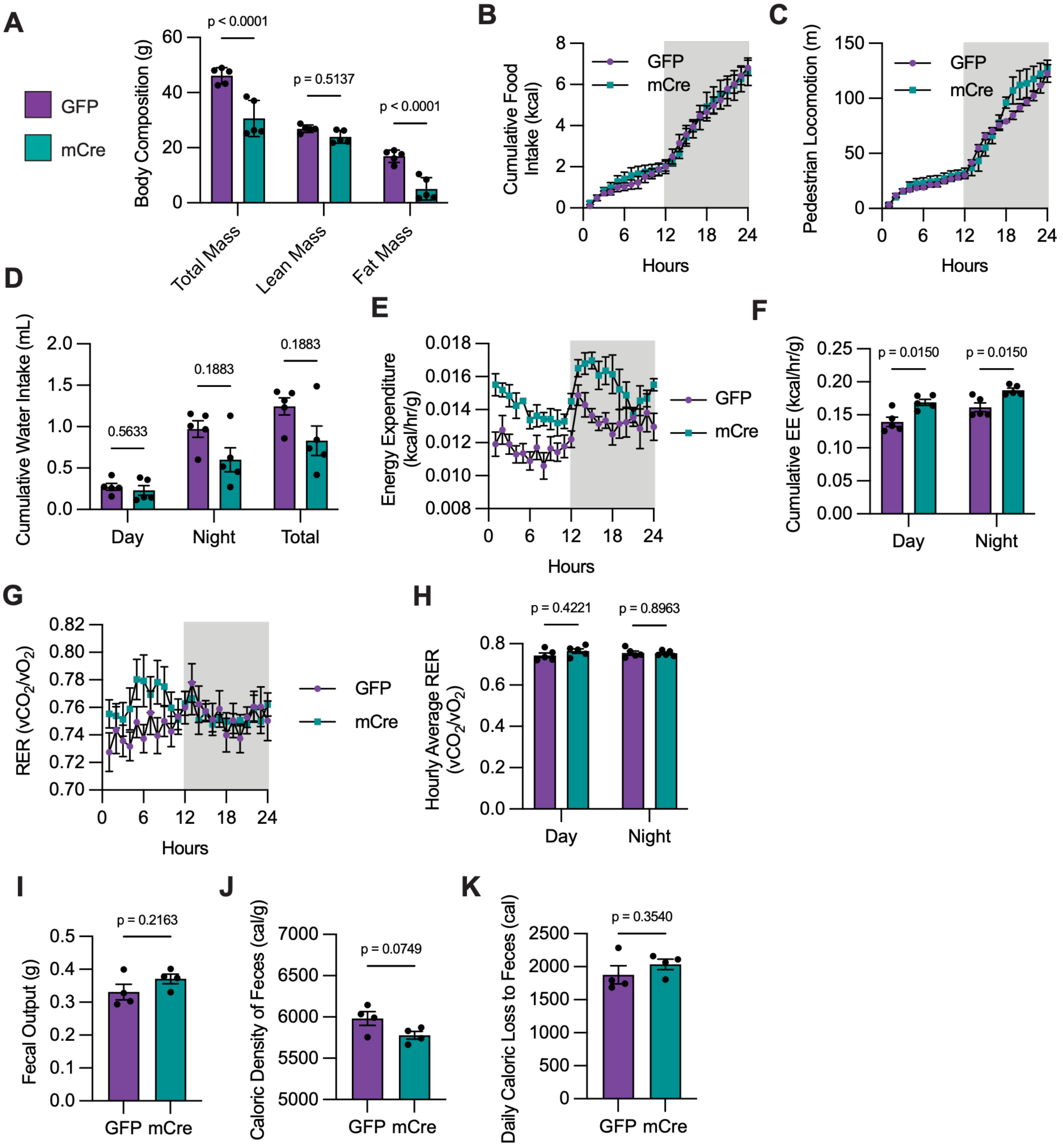


**Fig. S8. Mosaic deletion of hepatic *Ass1* increases energy expenditure.**

(**A-H**) AAV8-TBG-GFP control or -Cre (mCre, 2x$10^{10}$ vg/mouse) was administered to *Ass1*$^{fl/fl}$ mice at 6 weeks of age. HFD feeding was started at the time of AAV delivery. After 20 weeks, animals were moved to Promethion metabolic cages for indirect calorimetry analysis and DEXA PIXImus for body composition analysis. N=5 per condition. (**A**) Total, lean, and fat mass as measured by DEXA PIXImus. (**B**) Cumulative food intake. (**C**) Pedestrian locomotion. (**D**) Cumulative water intake for light (Day) and dark (Night) cycles. (**E**) Energy expenditure. (**F**) Cumulative energy expenditure summed for the light (Day) and dark (Night) cycles. (**G**)

Respiratory exchange ratio, RER. (**H**) Cumulative RER summed for the light (Day) and dark (Night) cycles. (**I-K**) GFP (n=4) or mCre (n=4) animals fed HFD for 20 weeks were singly housed for 24 hours, and total fecal matter was collected for bomb calorimetry. (**I**) Mass of fecal output. (**J**) Caloric density of feces. (**K**) Caloric loss to feces.

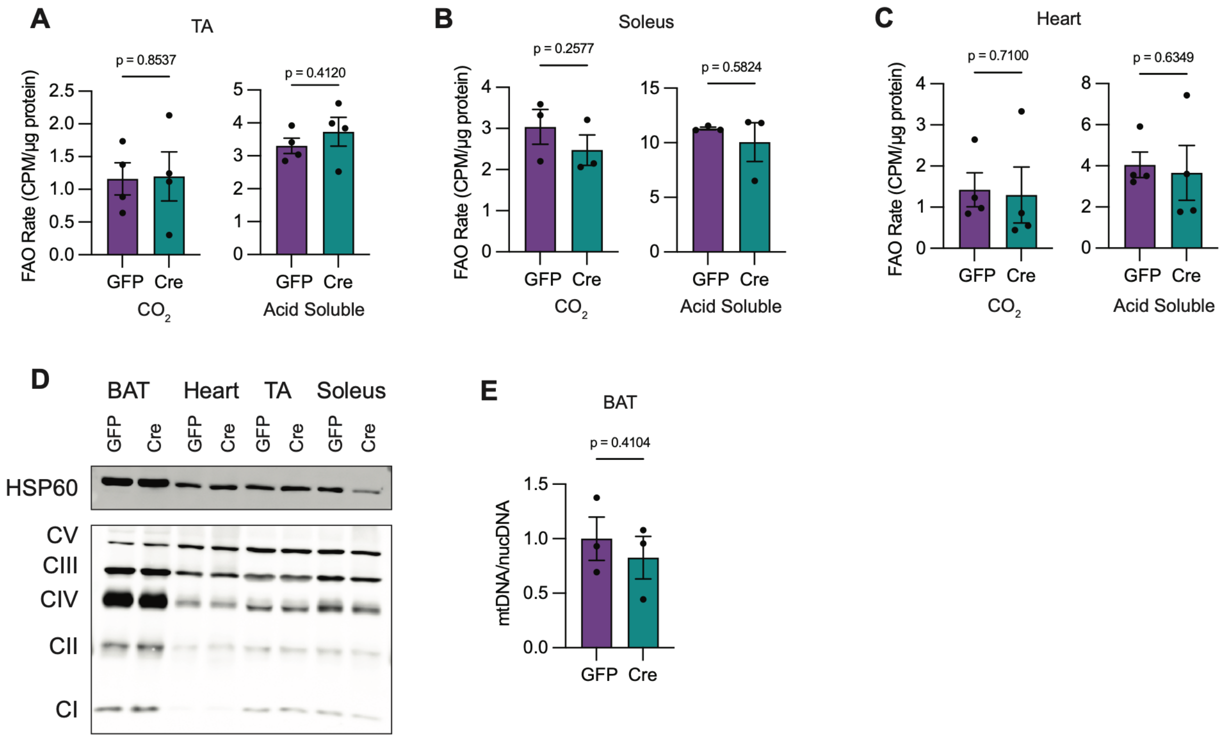


**Figure S9. FAO is unchanged in other oxidative tissues.**
(**A-C**) Mitochondria were isolated from TA (**A**), soleus (**B**), and heart (**C**) of GFP and Cre littermate pairs (n=3 or 4) fed HFD for 14 days. Fatty acid oxidation rates were assessed by $^{14}C$-Palmitate oxidation to $^{14}CO_2$ (complete oxidation) or $^{14}C$-acid soluble metabolites (incomplete oxidation). CPM, counts per minute. (**D**) Representative western blot of mitochondrial oxidative phosphorylation proteins in multiple oxidative tissues from a GFP and Cre littermate pair fed HFD for 14 days. (E) Mitochondrial DNA (mtDNA) to nuclear DNA (nucDNA) ratio in BAT tissue from GFP and Cre animals fed HFD for 14 days (n=3).

| Component | % kcal from | % by weight | Total kcal/g |
|---|---|---|---|
| *Standard Chow Diet (PicoLab Rodent Diet 20 #5053)* | | | |
| Protein | 24.495 | 21.0 | |
| Fat | 13.122 | 5.0 | |
| Carbohydrates | 62.382 | 53.5 | |
| | | | 4.11 |
| *High Fat Diet* | | | |
| Protein | 18.3 | 23.5 | |
| Fat | 60.3 | 27.3 | |
| Carbohydrates | 21.4 | 34.3 | |
| | | | 5.1 |
| *Gubra-Amylin NASH Diet* | | | |
| Protein | 20 | 22.5 | |
| `Fat | 40 | 19.9 | |
| Carbohydrates | 40 | 44.9 | |
| | | | 4.49 |
| *Ketogenic Diet* | | | |
| Protein | 20 | 31 | |
| Fat | 80 | 54 | |
| Carbohydrates | 0 | 0.2 | |
| | | | 6.1 |

**Table S1. Nutrient information of various diets**

| **SYBR Green** | | |
|---|---|---|
| **Gene** | **Forward** | **Reverse** |
| Ass1 (exon 12-13) | ATTGACATCGTGGAGAACCG | GTAAAGGATGGTCCCTGCTG |
| Actb | GGCTGTATTCCCCTCCATCG | CCAGTTGGTAACAATGCCATGT |

| **TaqMan** | |
|---|---|
| **Gene** | **Catalog Number** |
| Elovl2 | Mm00517086_m1 |
| Elovl3 | Mm00468164_m1 |
| Elovl5 | Mm00506717_m1 |
| Acsl5 | Mm01261083_m1 |
| Scd1 | Mm00772290_m1 |
| Scd2 | Mm01208542_m1 |
| Acaca | Mm01304258_m1 |

**Table S2. Primer Sequences for RT-qPCR**
